# *Wolbachia* feminizes a spider host with assistance from co-infecting symbionts

**DOI:** 10.1101/2025.02.13.638118

**Authors:** Virginija Mackevicius-Dubickaja, Yuval Gottlieb, Jennifer A. White, Matthew R. Doremus

**Author notes:** Data availability statement: All data associated with this manuscript are available in the supplemental material. Author contributions: All authors designed the experiments; VMD and MRD conducted the experiments; VMD and MRD conducted statistical analyses and generated figures; All authors contributed to the manuscript and approved the final version.

## Abstract

Arthropods commonly harbor maternally-transmitted bacterial endosymbionts that manipulate host biology. Multiple heritable symbionts can co-infect the same individual, allowing these host-restricted bacteria to engage in cooperation or conflict, which can ultimately affect host phenotype. The spider *Mermessus fradeorum* is variably infected with up to five heritable symbionts: *Rickettsiella* (R), *Tisiphia* (T), and three strains of *Wolbachia* (W1-3). Quintuply infected spiders are feminized, causing genetic males to develop as phenotypic females and produce almost exclusively female offspring. By comparing feminization across nine infection combinations, we identified a feminizing strain of *Wolbachia*, W1. We also observed that spiders infected with both W1 and W3 produced ∼10% more females than those lacking W3. This increase in feminization rate does not seem to be due to direct changes in W1 titer, nor does W1 titer correlate with feminization rate. Instead, we observed subtle titer interactions among symbionts, with lower relative abundance of R and T symbionts in strongly feminized infections. This synergistic effect of co-infection on *Wolbachia* feminization may help promote the spread of all five symbionts in spider populations. These results confirm the first instance of *Wolbachia*-induced feminization in spiders and demonstrate that co-infecting symbionts can improve the efficacy of symbiont-induced feminization.

## Introduction

Maternally-transmitted bacterial endosymbionts commonly infect terrestrial arthropods. These host-restricted heritable symbionts play major roles in the biology of their host, often manipulating host reproduction to favor infected females that transmit the symbiont (Doremus and Hunter, 2020). Heritable symbionts, such as the common symbiont *Wolbachia*, can use several different strategies to manipulate host reproduction ranging from inducing asexual reproduction, to causing mating incompatibilities between differentially infected hosts, or biasing host development to favor females (Doremus and Hunter 2020). Some heritable bacteria can also bias host development to favor females by feminizing genetic males, causing them to develop instead as phenotypic females capable of reproducing and transmitting the symbiont (Doremus and Hunter, 2020; Werren et al., 2008). *Wolbachia* are responsible for feminizing a diverse range of hosts including isopods (Bouchon et al., 1998), lepidopterans (Hiroki et al., 2002; Sugimoto and Ishikawa, 2012), and planthoppers (Negri et al., 2006). Feminization is also implicated in *Wolbachia*-induced parthenogenesis of some hymenopteran parasitoids (Fricke and Lindsey, 2024). Feminizing symbionts can alter the dynamics of host populations by causing female-biased sex ratios (Bourtzis and Miller, 2008; Hatcher et al., 1999; Kageyama et al., 2012), male rarity, and in extreme cases, may even influence the evolution of host sex determination systems (Bourtzis and Miller, 2008; Cordaux et al., 2011; Cordaux and Gilbert, 2017; Sugimoto and Ishikawa, 2012). Despite affecting a range of arthropod hosts, *Wolbachia* feminization remains comparatively less characterized compared to its other forms of reproductive manipulation ((Beckmann et al., 2017; LePage et al., 2017; Shropshire et al., 2018; Perlmutter et al., 2019; Fricke and Lindsey, 2024).

When multiple heritable symbiont strains co-infect the same host, this provides an opportunity for inter-strain interactions that can favor certain symbiont combinations over others (Doremus and Oliver, 2017; Leclair et al., 2017; Peng et al., 2023; Weldon et al., 2020; Zhu et al., 2012). These interactions include possible conflict over resources or access to intracellular niches within the host (Goto et al., 2006; Yang et al., 2020), or cooperation to facilitate their transmission (Peng et al., 2023). Symbiont co-infections can alter symbiont titer (Goto et al., 2006; Kondo et al., 2005; McLean et al., 2018; Yang et al., 2020), tissue tropism (Goto et al., 2006), transmission rate (Rock et al., 2018), and metabolic complementation (Peng et al., 2023), all of which can ultimately influence the penetrance of symbiont-induced phenotypes and the stability of symbiont co-infections.

Symbiont co-infections are particularly common in spiders, which often host heritable symbiont communities whose complexity rivals that of their better-studied insect cousins (Goodacre et al., 2006; Vanthournout and Hendrickx, 2015; Zhang et al., 2018; White et al., 2020). For example, North American populations of *Mermessus fradeorum*, a cosmopolitan dwarf spider, are variably infected with up to five heritable symbionts: *Rickettsiella, Tisiphia* (formerly *Rickettsia*; Davison et al., 2022), and three strains of *Wolbachia* (Curry et al., 2015; Rosenwald et al., 2020). The most common symbiont, *Rickettsiella*, usually infects >99% of spiders in a population and causes a form of reproductive sabotage called cytoplasmic incompatibility (Rosenwald et al., 2020). The next most common symbiont combination (symbiotype) is all five symbionts co-infecting the same spider; these quintuply infected spiders are feminized, but the symbiont(s) responsible for feminization have yet to be identified (Curry et al., 2015). The *M. fradeorum* system provides an excellent opportunity to study multiple forms of reproductive sabotage in the same host and to learn more about the biology of heritable symbioses of spiders (Goodacre et al., 2006; Vanthournout and Hendrickx, 2015; Zhang et al., 2018; White et al., 2020) but further research requires identifying the symbiont(s) responsible for feminization.

By comparing offspring sex ratios among spiders harboring nine different symbiotypes, we identified a strain of *Wolbachia* (W1) as necessary for feminization in *M. fradeorum*. Co-infection with certain symbionts synergistically improved W1 feminization, which may reinforce co-transmission of the five-member symbiont community in nature. We also found that W1 numerically dominates the heritable symbiont community and is consistently the most abundant symbiont when present. Despite the overall higher titer of W1 compared to other symbionts in this system, W1 titer was not associated with feminization rate. Instead, the relative abundance of R and T symbionts was reduced in strongly feminized symbiotypes, suggesting that improved *Wolbachia* feminization may arise from subtle interactions among co-infecting symbionts. These results confirm the first instance of *Wolbachia*-induced feminization in a spider, inform our understanding of *M. fradeorum* feminization, and demonstrate that symbiont co-infections can have synergistic effects that increase the penetrance of symbiont-conferred feminization.

### Experimental Procedures

The *M. fradeorum* cultures used in this study were initially collected from alfalfa fields in Kentucky, USA, and subsequently maintained in the laboratory. Spiders were kept at 20 °C in individual 4 cm diameter rearing cups with moistened plaster at the bottom for humidity control (Rosenwald et al., 2020). Immature spiders were fed collembola (*Sinella curviseta*) until they neared adulthood; thereafter they were fed one wingless *Drosophila melanogaster* twice a week.

North American field populations of *M. fradeorum* are naturally infected with up to five heritable symbionts: *Rickettsiella* (R), *Tisiphia* (T), and three strains of *Wolbachia* (W1, W2, W3; Rosenwald et al., 2020). For simplicity we will refer to symbionts by their letter abbreviations. To identify which of these five symbionts contribute to feminization, we used a panel of 9 symbiotypes including quintuply infected (RTW123), quadruply infected (RW123, RTW12, RTW23), triply infected (RTW1, RTW2), doubly infected (RT), singly infected (R), and uninfected spiders. All infected symbiotypes included *Rickettsiella* because we were unable to generate or maintain lineages that lost *Rickettsiella* without losing the other symbionts. These combinations included both naturally occurring symbiotypes and experimental symbiotypes generated via past antibiotic administration (Curry et al., 2015; Rosenwald et al., 2020). Antibiotic treatments consisted of misting developing spiders with a fine spray of tetracycline (0.1%) and ampicillin (0.1%) until sub-adulthood (Rosenwald et al., 2020). All lines produced via antibiotic treatments were maintained in the lab for at least 5 generations prior to use in experiments.

We mated 8–12-week-old female spiders representing the 9 symbiotypes with uninfected male spiders. After females laid one egg mass, they were held for five days in a 1.5 mL microcentrifuge tube to void their gut contents and then were stored in 95% ethanol at -20 °C until we extracted their DNA using DNeasy Blood & Tissue kits (Qiagen) following the manufacturer’s protocol. To estimate sex ratio, we randomly separated up to 8 siblings per egg mass into individual 4 cm cups to avoid cannibalism. Egg masses that produced fewer than 4 offspring were removed from the experiment. These offspring were reared at 20 °C and were first provided *S. curviseta* and then *D. melanogaster* as they developed. We determined offspring sex morphologically by the presence of enlarged pedipalps in males once spiders reached the penultimate immature stage. Using logistic regression with a quasibinomial distribution (R v 4.4.0, R Team), we tested the effect of symbiont infection on spider sex ratio using a series of planned contrasts between infected spiders and uninfected control spiders. These contrasts include a full model comparing the entire set of eight infected symbiotypes and the uninfected control, as well as contrasts between each individual symbiotype and the control. We next used logistic regressions to compare feminization rates between each feminized symbiotype and the quintuply infected line to see if the presence or absence of co-infecting symbionts modified the level of feminization.

We also assessed whether symbiont titer was related to feminization. We conducted digital PCR (dPCR) assays to quantify symbiont genome copies in feminized *M. fradeorum* samples using short specific primers and probes designed for symbiont and spider genes as specified in **Table 1**. We used Geneious Prime Version 2021.2 software (https://www.geneious.com/updates) to design primers and probes, DINAMelt Web Server (http://www.unafold.org/Dinamelt/applications/two-state-melting-folding.php) to predict hybridization and folding of the amplicons, and BLAST Web Server (https://blast.ncbi.nlm.nih.gov/Blast.cgi) to test primer specificity. Primers and probes were obtained from Biolegio BV (Nijmegen, Netherlands) and were dissolved in TE buffer (Tris-EDTA; 10 mM Tris base, 0.1 mM EDTA, pH 8.0) to make 100 μM stocks and further diluted with TE buffer to create 20 μM working solutions (stocks and working solutions stored at -20 °C until use). TaqMan probes were synthesized by labeling the 5’ terminal nucleotide with a fluorophore and the 3’ terminal nucleotide with a dark quencher, Black Hole Quencher 1 (BHQ1) or Black Hole Quencher 2 (BHQ2). Probes and primers were tested for specificity using diagnostic PCR and quantitative PCR (diagnostic and qPCR details included in Supplemental material).

**Table 1.**
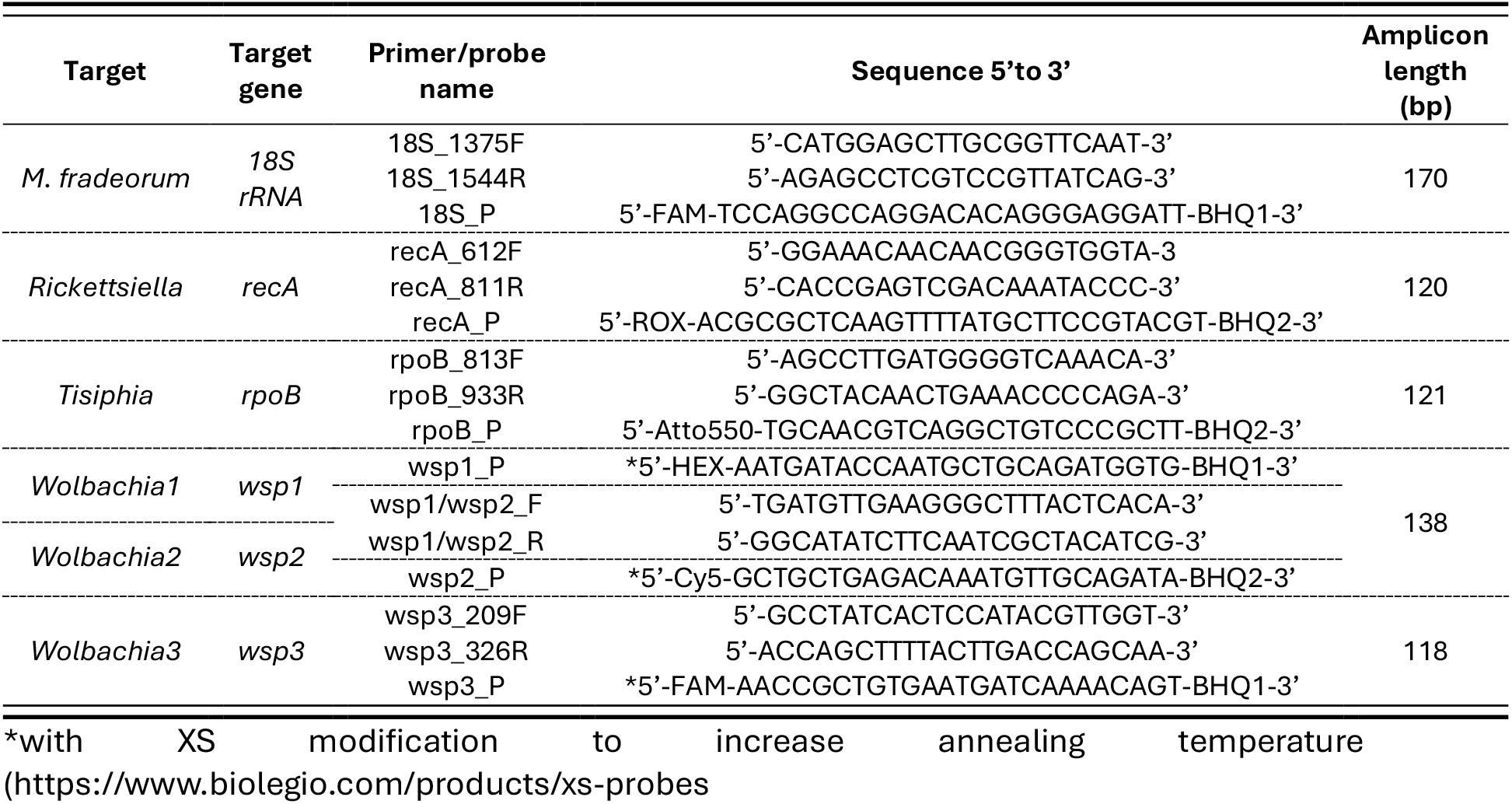
Primers and probes used for digital PCR.

We performed the detection and amplification of the spider *18S rRNA* gene separately from symbiont genes using Qiagen QIAcuity Probe PCR Kit and QIAcuity One-plate digital PCR instrument (QIAcuity One, 5plex instrument, Qiagen, Germany). The dPCR mixture for a single reaction was 10 μL of 4x Probe QIAcuity PCR Master Mix, 0.8 μM final concentration of each primer, 0.4 μM final concentration of probe, 12 ng / 1.2 ng of samples gDNA for symbiont / 18S rRNA gene, respectively, and RNase-free water for a total reaction volume of 40 μL. Once prepared, the reaction mixtures were transferred to QIAucity Nanoplate 26k 24-well and sealed with the QIAcuity Nanoplate Seal. We used the following dPCR cycling program: 2 min at 95 °C for initial heat activation, followed by 40 cycles of 95°C for 15 s for denaturation, and 60 °C for 30 s for annealing and extension. All dPCR runs contained negative controls containing the same DNA background without the targeted sequence and/or DNA Elution buffer AE (10 mM Tris-HCL, 0.5 mM EDTA, pH 9, Qiagen), and the same two quintuply infected positive controls to enable data comparison across multiple nanoplates. After the reaction finished, the signal was quantified, and we set thresholds manually to separate two populations: positive partitions with increased fluorescence intensity and negative partitions with baseline signal only. Reactions containing below 100 random unexpected positive partitions of the symbiont gene with a high confidence interval (CI) were considered negative for that symbiont. We calculated gene copy number per μL for each gene using the integrated QIAcuity Software Suite. To adjust for the gDNA input amount (12 ng of sample for symbiont detection,1.2 ng of sample for spider gene detection for each dPCR reaction), we multiplied *18S rRNA* copies/μL by this gDNA dilution factor. Normalized symbiont titers (*normalized symbiont titer = symbiont gene copy number/ 18S rRNA gene copy number)* for every symbiont were compared across feminized symbiotypes using Kruskal-Wallis tests, followed by pairwise Mann-Whitney U-Tests to determine differences between specific pairs of symbiotypes in R (R v 4.4.0, R Team). We also tested for a correlation between W1 titer and feminization rate using Spearman’s correlation test in R (R v 4.4.0, R Team).

The initial crossing experiment showed an effect of symbiont co-infection on feminization rate, with spiders infected with all five symbionts showing increased feminization rates compared to spiders infected with subsets of fewer symbionts. To confirm that the observed differences in feminization between RTW123 and RTW1 spiders were replicable, we performed an additional experiment comparing the sex ratio among uninfected (n = 12), triply infected (RTW1; n = 12), and quintuply infected (RTW123; n = 12) spiders. As in the previous experiment, eight-week-old female spiders were mated with uninfected males, allowed to lay one egg mass, starved for 5 days then stored in 95% ethanol at -20 °C prior to DNA extraction. For this set, symbiont infection status was confirmed with diagnostic PCR (supplemental material). We estimated feminization rates by morphologically characterizing the sex of 9 randomly selected offspring as in the first assay. We then compared feminization rates across these three infection types using logistic regression with a quasibinomial distribution in R (R v 4.4.0, R Team).

## Results

### Co-infection modifies the efficacy of Wolbachia 1-induced feminization

Offspring sex ratio (**Fig. 1**) varied significantly across spiders harboring different symbiotypes (ΔDeviance = 189.22, d.f. = 8, p < 0.001). Compared to uninfected spiders, which generally produce an even sex ratio, quintuply infected female spiders (RTW123) produced almost exclusively female offspring as expected (ΔDeviance = 36.34, d.f. = 1, p < 0.001; Curry et al 2015). Other symbiotypes that contained W1 (RTW1, RTW12, RW123) also exhibited strongly female-biased sex ratios compared to uninfected spiders (ΔDeviance_RTW1_ = 24.89, d.f. = 1, p < 0.001; ΔDeviance_RTW12_ = 20.61, d.f. = 1, p = 0.001; ΔDeviance_RW123_ = 30.82, d.f. = 1, p < 0.001). Spiders lacking W1 (R, RT, RTW2) did not differ from uninfected controls (ΔDeviance_R_ = 0.64, d.f. = 1, p = 0.345; ΔDeviance_RT_ = 0.65, d.f. = 1, p = 0.404; ΔDeviance_RTW2_ = 0.06, d.f. = 1, p = 0.768). Strikingly, spiders infected with all symbionts except W1 also were not feminized compared to uninfected spiders (ΔDeviance_RTW23_ = 0.14, d.f. = 1, p = 0.607), confirming that W1 is essential for feminization and is likely the sole inducer of this phenotype in *M. fradeorum*.

**Figure 1.**
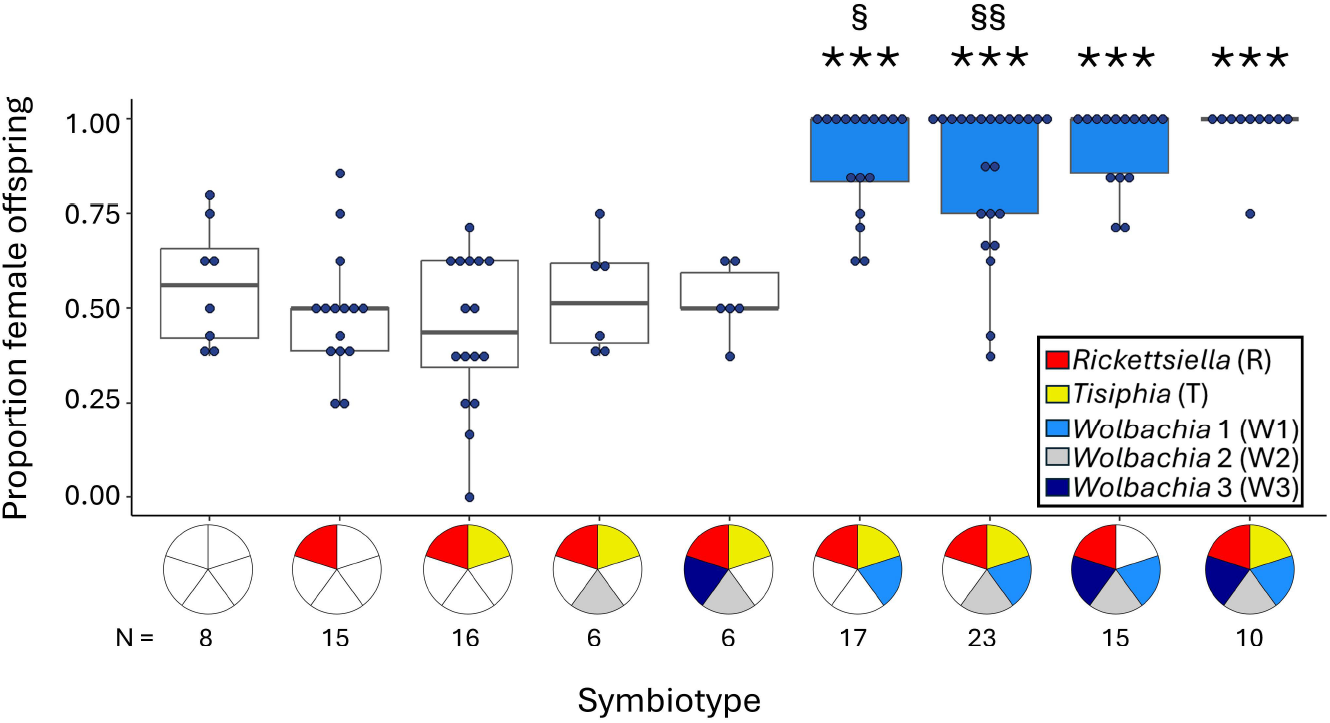
Proportion of female offspring produced by female *M. fradeorum* infected with different symbiont combinations. Symbiotypes are represented with colored pie charts, with each portion of the chart representing a symbiont as indicated in the key. Significant differences in proportion of female offspring compared to uninfected females are denoted with asterisks (*** = p <0.001). Significant differences in proportion female offspring of different W1 infection combinations compared to RTW123 spiders are denoted by the symbol § (§ = p <0.05, §§ = p <0.01). Box colors represent W1 infection, with infections lacking W1 colored white and infections including W1 colored blue. Sample sizes are listed below infection status.

However, not all combinations containing W1 were equal. Quintuply infected spiders produced a higher proportion of female offspring than either RTW1 (ΔDeviance = 4.46, d.f. = 1, p = 0.03) or RTW12 spiders (ΔDeviance = 8.43, d.f. = 1, p = 0.004). Feminization rates of RW123 spiders were similar to those of quintuply infected spiders (ΔDeviance = 1.56, d.f. = 1, p = 0.212), indicating that certain combinations of co-infecting symbionts can influence the penetrance of W1 feminization. When W1-infected spiders also harbored W3 (RW123, RTW123), their feminization rates were ∼10% higher than those that lacked this additional *Wolbachia* strain.

A second, more targeted assay compared feminization rates among uninfected, RTW1, and RTW123 spiders also found stronger feminization in RTW123 than RTW1 spiders (**SI Fig. 1**; ΔDeviance = 8.01, d.f. = 1, p = 0.029). As in the first experiment, feminization was ∼10% stronger in RTW123 spiders than RTW1 spiders, reinforcing the finding that co-infection by other symbionts strengthens feminization.

### Symbiont titers show limited change across infection types and feminization rates

In the main experiment, normalized W1 titers only marginally changed across the four feminized symbiotypes (K-W χ^2^ = 6.38, d.f. = 3, p-value = 0.09). Pairwise comparisons showed W1 titer to be lower in RTW1 spiders than RTW123 spiders (Mann-Whitney U-test p-value = 0.036; **Fig. 2**), with other symbiotype combinations having intermediate W1 titers (MWU p-value = 0.278-0.870 for all comparisons). *Rickettsiella* titer showed a significant change across symbiotypes (Kruskal-Wallis χ^2^ = 7.98, d.f. = 3, p-value = 0.05), possibly driven by reduced R titer in RW123 spiders (**Fig. 2**). Yet pairwise comparisons amongst R-containing symbiotypes did not show significant differences (MWU p-value = 0.092-0.623 for all comparisons). The titers for T (K-W χ^2^ = 0.79, d.f. = 2, p-value = 0.67), W2 (K-W χ^2^ = 3.07, d.f. = 2, p-value = 0.21), and W3 (K-W χ^2^ = 0.01, d.f. = 1, p-value = 0.91) did not change across their respective infected symbiotypes (**Fig. 2**). A Spearman correlation test did not support a correlation between sex ratio and normalized W1 titer (**SI Fig. 2**; Spearman’s *r*_*s*_ = 0.026, p-value = 0.84).

**Figure 2.**
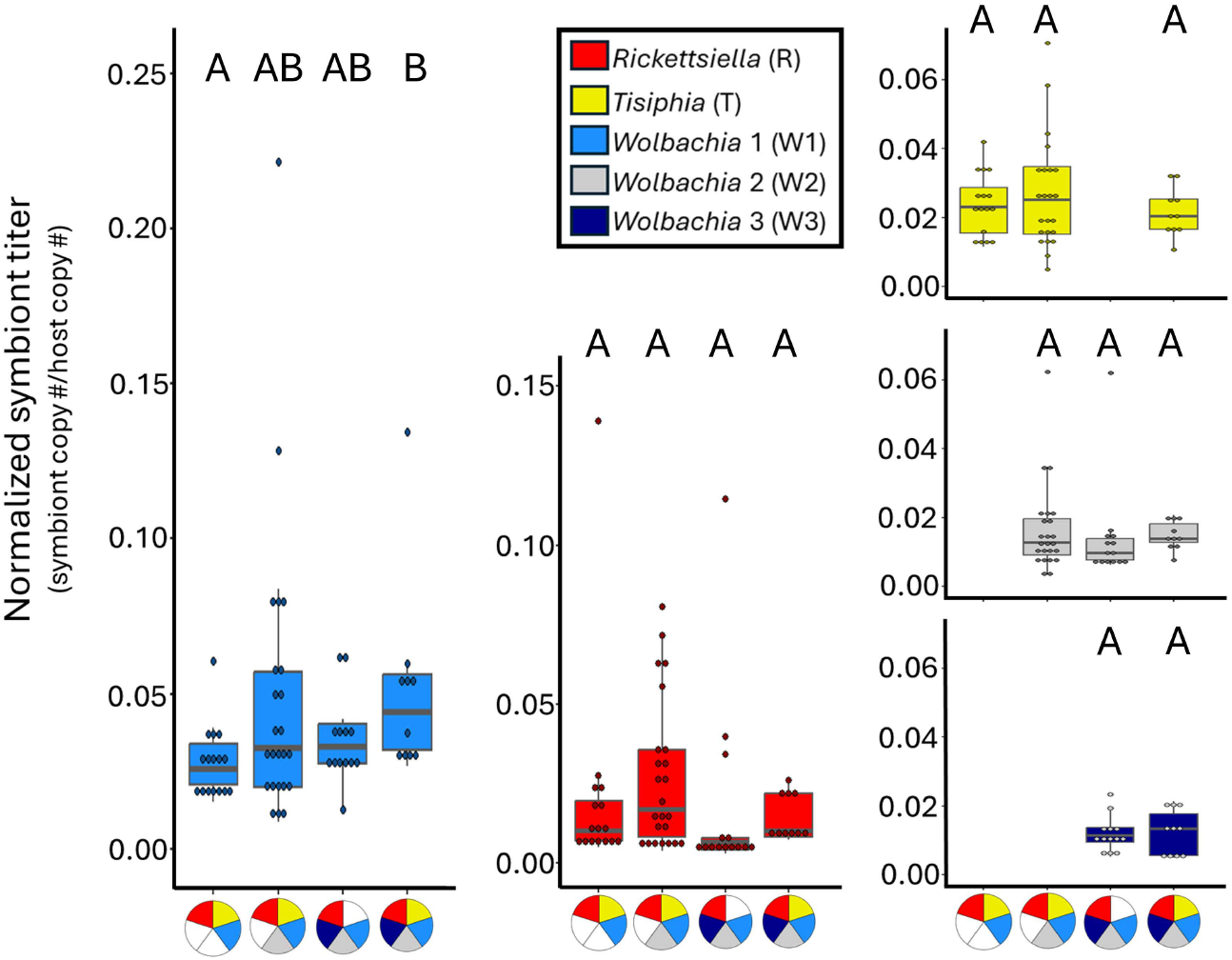
Symbiont titers estimated with dPCR normalized to host 18s rRNA in spiders infected with feminizing symbiotypes. Inset key indicates which symbiont is depicted in which panel and which symbiotypes are represented by pie charts on the x-axes. Symbiont genes used in dPCR were *recA* (*Rickettsiella*), *rpoB* (*Tisiphia*), and *wsp* (*Wolbachia 1-3*). Normalized symbiont titer was compared between infection types using multiple pairwise Mann-Whitney U-tests with Benjamini-Hochberg corrected p-values. Letters represent significantly different normalized titers (p < 0.05).

### Wolbachia 1 *relative abundance is highest and stable across feminized infection types*

Despite relatively subtle changes in symbiont titer, we did observe qualitative changes in the relative abundance of some symbionts symbiotypes (**Fig. 3**). The relative abundance of each symbiont is expected to fall as more symbionts are added to the community. For example, W3 is relatively more abundant in quadruple infections (RW123, RTW23) compared to the quintuple infection (**Fig. 3**). Yet the abundance of W1 relative to other symbionts remained stable across infections, with W1 comprising ∼40 - 50% of the total symbiont titer. Other symbionts, most notably R and T, exhibited a marked reduction in relative abundance in the most feminized symbiotypes, RW123 and RTW123.

**Figure 3.**
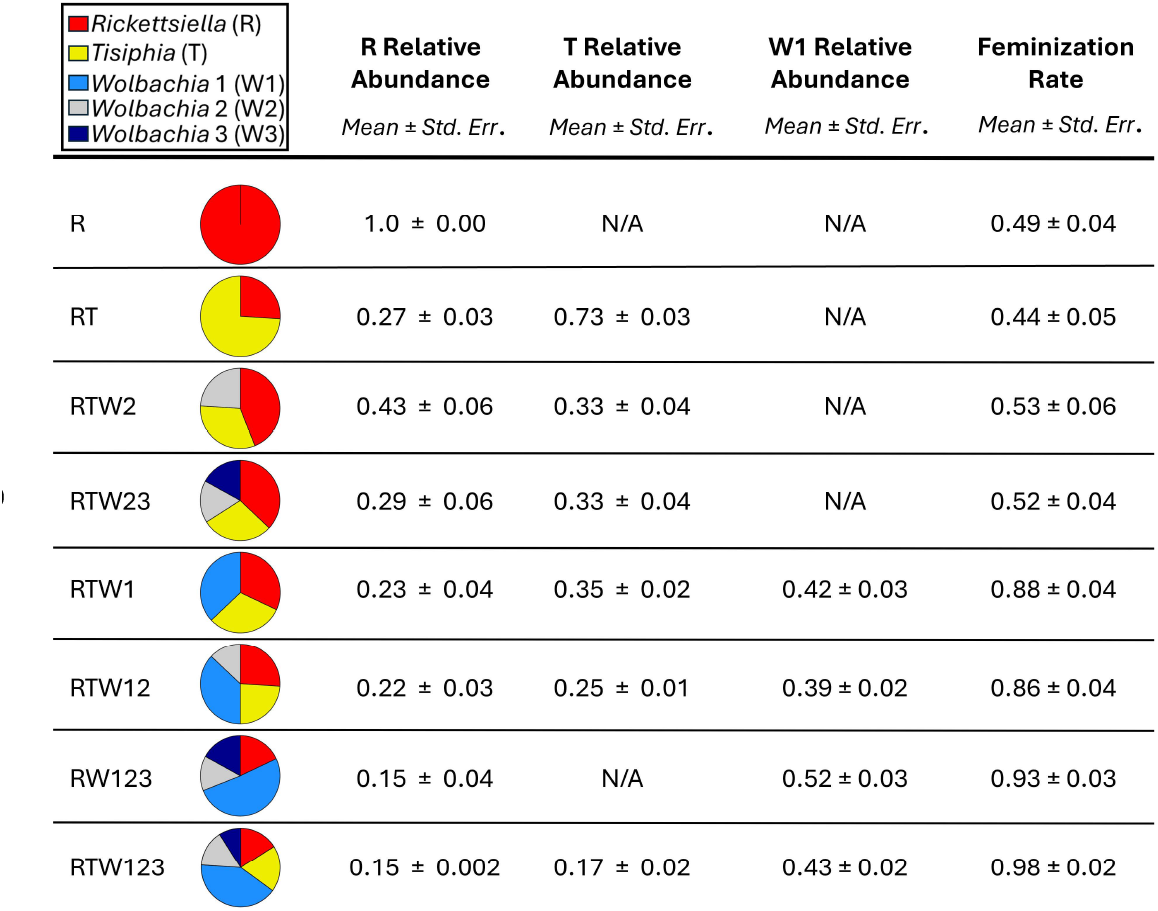
Mean relative abundance for R, T, and W1 in spiders with different infection combinations and feminization rates. The relative abundance of symbionts is expected to fall as the number of symbiont strains increases, but *Wolbachia* 1 maintains the highest proportion regardless of other members of the symbiont community. R and T represent the highest proportion when *Wolbachia* 1 is not present, with their lowest relative abundances occurring in symbiotypes with the highest feminization rates.

*Rickettsiella* comprises ∼30-40% of the symbiont titer in co-infected symbiotypes lacking W1, but its proportional representation drops to <20% of the total titer in RW123 and RTW123 spiders. Those symbiotypes also featured a similar reduction in T abundance, dropping from ∼30% to <20% of total symbiont titer. These changes suggest that the structure of this heritable symbiont community undergoes a substantial shift favoring *Wolbachia* 1 when this symbiont is present.

## Discussion

We found that one *Wolbachia* symbiont (W1) is required for feminization in *M. fradeorum* (**Fig. 1)**. Spiders harboring W1 exhibited female-biased sex ratios, while those lacking W1 did not. As we did not evaluate the phenotype of spiders singly infected with W1, it remains possible that *Rickettsiella* (and/or other symbionts like *Tisphia*) might also play a direct role in feminization when W1 is present. Despite numerous attempts, we have been unable to generate symbiotypes that do not include *Rickettsiella* in this system. Our results therefore highlight that W1 is a necessary component for feminization of *M. fradeorum*, but do not yet show that it is sufficient for feminization on its own.

We also found a synergistic effect of co-infection on *Wolbachia* feminization, with spiders harboring two symbiotypes, RTW123 and RW123, exhibiting ∼10% stronger feminization than other feminized symbiont combinations (**Fig. 1, SI Fig. 1**). This synergistic increase in feminization may involve direct changes in the feminizing *Wolbachia* titer, as both W1 titer and feminization rate were increased in RTW123 spiders relative to the more variably feminized RTW1 spiders (**Fig. 2**). However, because feminization rates were generally high in this system across all symbiotypes bearing W1, we ultimately did not find a correlation between W1 titer in adult females and their feminization rate (**SI Fig. 2**).

Symbiont titer at other stages of development (e.g. embryogenesis) might also be more reflective of realized feminization rates (Doremus et al., 2019; Herran et al., 2020; Doremus et al., 2022). Additionally, while *Wolbachia* titer is generally thought to correspond with feminizing phenotypic penetrance (Rigaud et al., 2001; Narita et al., 2007; Herran et al., 2020), there are instances of low-titer *Wolbachia* capable of inducing strong manipulative phenotypes (Richardson et al., 2019). It is possible that W1 only requires a low titer threshold for reliable feminization induction, which would further limit our ability to identify a correlation between titer and phenotype expression.

The influence of titer on feminization is seemingly subtle in this system and could also involve changes in the composition of the entire symbiont community (**Fig. 3**), rather than the normalized titer of any one symbiont. The composition of the symbiont community shifts in feminized spiders to favor W1, which seems to dominate the symbiont community when present. This shift comes largely at the expense of the relative abundance of *Rickettsiella* and *Tisiphia*, with the remaining *Wolbachia* strains showing more limited changes in relative abundance across symbiotypes (**Fig. 3**). The variation in symbiont community composition hints at potential competitive interactions among symbionts that seemingly favor *Wolbachia* 1.

This web of symbiont interactions may alter W1 biology in other ways beyond shifting its titer, such as influencing W1 localization or gene expression. Many bacteria regulate their gene expression through quorum sensing in response to bacterial cell density, including that of other bacterial strains and species (Miller and Bassler, 2001). While research on quorum sensing in *Wolbachia* is limited, manipulation of the quorum sensing pathway has been implicated in affecting *Wolbachia* titer, as well as its CI and male-killing phenotypes (Hidayanti et al., 2022, 2023). If W1 uses quorum sensing to control elements of its gene expression, the presence of W3 and/or other symbionts could directly influence expression of feminization factors. Hypothetically, co-infecting symbionts could also indirectly affect feminization by altering host biology, perhaps by altering host developmental time during critical life stages or by producing co-factors that improve the efficacy of W1-encoded feminizing factors.

The feminizing capabilities of W1 should allow it to spread through spider populations regardless of its co-infecting partners, yet W1 is usually observed in quintuple infections or alongside W3 in nature (Rosenwald et al. 2020). This infection pattern is suggestive of a synergistic link between these symbionts (Rock et al., 2018; Rosenwald et al 2020; Peng et al. 2023). The augmentation of W1 feminization by co-infecting symbionts, particularly W3 (**Fig. 1, SI Fig. 1**), may promote the transmission of the quintuple infection. This cooperative effect clearly benefits symbionts like W3, which do not seem to directly influence host reproduction and thus benefit from hitchhiking off improved W1 feminization. However, it is not fully clear how this interaction benefits W1, as W1 is still able to induce feminization without W3, albeit with slightly reduced efficacy. The quintuple co-infection might additionally benefit from other synergistic effects, such as increased transmission rates, reduced infection costs of W1, and/or context-dependent benefits like protection from natural enemies (Rock et al., 2018). Assays comparing symbiotype effects on host and symbiont fitness may reveal additional synergistic effects of co-infection that reinforce the maintenance of the quintuple infection in nature.

Despite their prevalence, interactions among cohabiting symbionts remain understudied. The prominence of certain symbiotypes in host populations suggests that some symbiont combinations provide a selective advantage over others that can reinforce their shared transmission and spread (Peng et al. 2023). The effect of symbiotype on *Wolbachia*-induced feminization in *M. fradeorum* provides an example of cooperation among symbionts. Identification of this effect, and the characterization of the feminizing symbiont of *M. fradeorum*, are essential steps in understanding the dynamics that shape the heritable symbiont communities of arthropods.

## Acknowledgements

We thank K. Butler, A. Fajardo, V. Kegley, J. Proctor, E. Roberson, and E. Williams for assistance in the lab. We also thank E. Klement for discussions about statistical analyses.

## Notes

Funding statement: This material is based upon work supported by the National Science Foundation under Grant No. 1953223 and the Binational, USA-Israel, Science Foundation (BSF) under Grant No. 201697 to JAW and YG; the National Institute of Food and Agriculture, U.S. Department of Agriculture under Hatch Nos. 1020740 and 7007679 and Fellowship No. 2023-67012-39352 to MRD.

Conflict of interest disclosure: The authors declare no conflicts of interest.

### Competing Interest Statement

The authors have declared no competing interest.

