## Supplemental Material for "*Wolbachia* feminizes a spider host with assistance from co-infecting symbionts"

*Confirmation of dPCR probe and primer specificity*

We confirmed the specificity of the dPCR probe and primer pairs with qPCR. The qPCR mixture for a single reaction was 10 μL of 4x Probe QIAcuity PCR Master Mix, 0.8 μM final concentration of each primer, 0.4 μM final concentration of probe, 30 ng of samples gDNA, and RNase-free water for a total reaction volume of 40 μL. Once prepared, the reaction mixture was run using the Step One Plus real-time PCR system (Applied Biosystems, Ca, USA). We used the following qPCR cycling program: 2 min at 95 °C for initial heat activation, followed by 40 cycles of 95 °C for 15 s for denaturation, and 60 °C for 30 s for annealing and extension. All qPCR runs were performed in triplets and contained negative controls containing the same DNA background without the targeted sequence and/or DNA Elution buffer AE (10 mM Tris-HCL, 0.5 mM EDTA, Ph 9, Qiagen).

Primers used in diagnostic PCR

To confirm symbiont infections of spiders used in the second mating assay , we first extracted DNA using the DNeasy Blood and Tissue kit (Qiagen) following the manufacturer’s protocol. We then performed diagnostic polymerase chain reactions (PCR) for *Rickettsiella*, *Tisiphia*, and three *Wolbachia* strains on females to confirm their infection status. For *Rickettsiella* and *Tisiphia*, we used previously published diagnostic primers (Curry et al., 2015; Duron et al., 2016; **SI Table 1**). We designed short primers specific for the *wsp* gene for each *Wolbachia* strain using Geneious Prime Version 2021.2 (**SI Table 1**). We tested the specificity of the wsp primers *in silico* and by using diagnostic PCR containing the same DNA background with/without the targeted sequence. The PCR mixture for a single reaction was 10 µL of GoTaq® Green Master Mix, 0.8 μM final concentration of each primer, 30ng of sample gDNA, and RNase-free water for a total reaction volume of 20 μL. Diagnostic PCRs were run using the X^96^ Thermal Cycler from VWR with the following cycling program: 3 min at 95 °C for initial denaturation, followed by 35 cycles of 30 s at 95 °C for denaturation, 24 s at 60 °C for annealing, and 1 min at 72 °C for extension, followed by a final extension at 72 °C for 10 min. Samples were electrophoresed on 1.5% agarose gels containing GelRed® Nucleic Acid Gel Stain (Biotium™). Quick Load 100bp DNA Ladder (New England Biolabs®) was run alongside samples at 80V for 45 min.

**SI Table 1** Primer pairs used in this study, with references provided for previously published primers for each symbiont used in diagnostic PCR.

| **Target** | **Target gene** | **Primer name** | **Sequence** | **Annealing Temp (°C)** | **Amplicon length (bp)** | **Citation** |
| --- | --- | --- | --- | --- | --- | --- |
| *Rickettsiella* | 16S rRNA | RLA16s F1 | CAGTAAARRTTTCGGYCTTTAYGGG | 56 | 532 | Duron et al. 2015 |
|  |  | RLA16s R1 | CGTGTAGGTGGTTGACTAGGTTTG |  |  |  |
| *Tisiphia* | 16S rRNA | RicklongF | ACGTGGGAATCTACCCATCA | 60 | 530 | Curry et al. 2015 |
|  |  | RicklongR | TAGCCTAGATGACCGCCTTC |  |  |  |
| *Wolbachia* 1 | wsp | wsp1_36F | ACAAAAGCATCAGGTCAAGAAAAT | 60 | 167 | This study |
|  |  | wsp1_201R | CATCTGCAGCATTGGTATCATTT |  |  |  |
| *Wolbachia* 2 | wsp | wsp2_93F | GCAAGGCAACAAATAAAGACAAGG | 60 | 163 | This study |
|  |  | wsp2_255R | ACATTTGTCTCAGCAGCAGC |  |  |  |
| *Wolbachia* 3 | wsp | wsp3_270F | ACCGCTGTGAATGATCAAAACA | 60 | 162 | This study |
|  |  | wsp3_431R | ATCGTTATTAGTTGATGTTGTTGCTT |  |  |  |

**SI Figure 1** A replicated mating trial comparing the proportion of female offspring produced by two *Wolbachia* 1 infection combinations (RTW1, RTW123) to uninfected spiders. Proportion of female offspring was compared across the three symbiotypes using logistic regression. Significant differences in female offspring production are denoted with different letters. Both RTW123 (ΔDeviance = 50.32, d.f. = 1, p <0.001) and RTW1 spiders (ΔDeviance = 19.14, d.f. = 1, p <0.001) were feminized relative to uninfected controls, but RTW123 spiders were more feminized than RTW1 spiders (ΔDeviance = 8.01, d.f. = 1, p = 0.029). Co-infection of the entire symbiont community again increased W1 feminization rate by ~10% (**Fig. 1).** Box colors represent W1 infection, with infections lacking W1 colored white and infections including W1 colored blue. Symbiotypes are represented with colored pie charts. Sample sizes are listed below infection status.

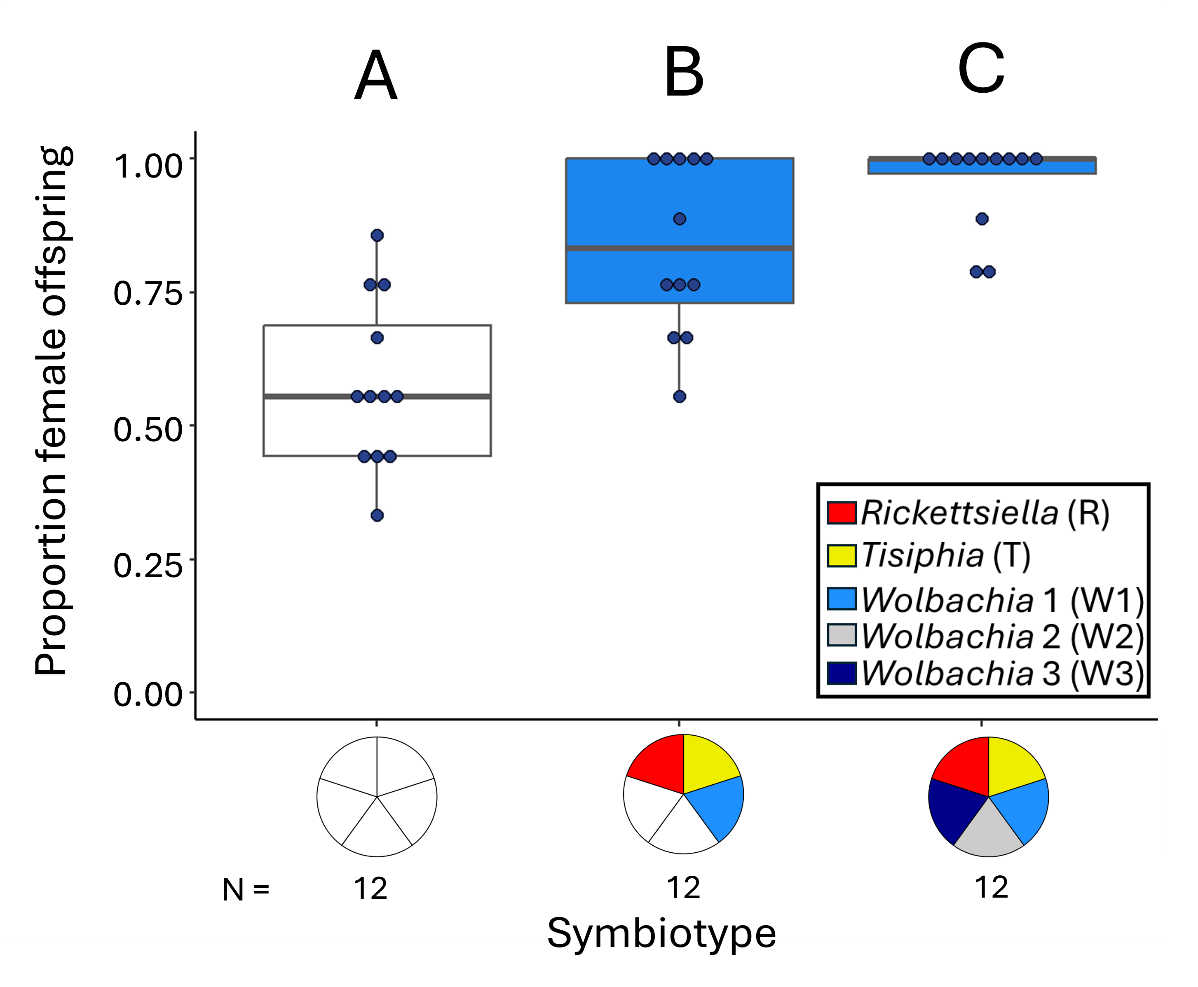

**SI Figure 2** Normalized Wolbachia 1 titer does not correlate with feminization rate. Shape and color of dots correspond with different symbiotypes. Correlation coefficients for the Spearman’s test is provided, p-value = 0.84.

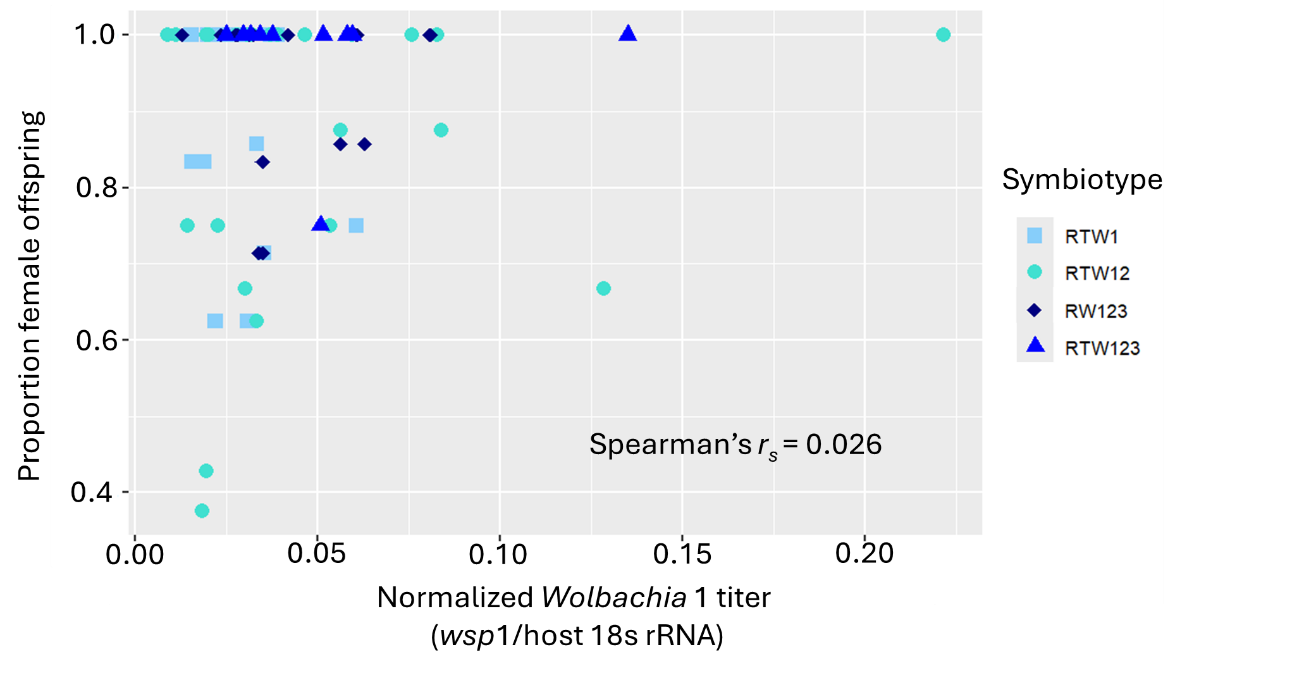
